# Sex-divergent trajectories of hippocampal and cortical NMDA receptor density across the Alzheimer’s disease continuum

**DOI:** 10.64898/2026.08.05.743051

**Authors:** Maricedes Acosta-Martínez, Vanessa Carter, Aviram Nessim, Sarah Murphy, Jasbeer Dhawan, Thomas G. Beach, Geidy E. Serrano, Erin E. Sundermann, Anat Biegon

## Abstract

While loss of NMDA receptors (NMDARs) is associated with Alzheimer’s disease (AD) severity, the effect of sex or the relationship between regional NMDAR density and antemortem cognitive status across the AD spectrum has not been examined. We performed quantitative *in vitro* autoradiography of hippocampus, entorhinal cortex (EC), and parietal cortex using NMDAR and tau radioligands. Relationships between regional NMDAR density and cognitive status assessed by the Mini Mental State Exam (MMSE), and between NMDAR and tau density, were examined by bivariate correlations. In both sexes, the largest AD-related decreases in NMDAR density were observed in the CA1 field. However, there was a significant diagnosis by sex interaction driven by sex-specific changes in the mild cognitive impairment (MCI) stage, with lower NMDAR density in MCI women, but not MCI men relative to same-sex controls. Within diagnosis analyses revealed positive correlations between NMDAR density and MMSE scores and significant negative correlations between EC NMDAR and tau density, which was significant only in AD men. Our data show that changes in hippocampal NMDAR density across the AD continuum are modulated by sex and may contribute to the known sex differences in the clinical trajectory of the disease.

## 1. Introduction

Alzheimer’s disease (AD) is a progressive neurodegenerative disorder characterized by deposition of amyloid plaques and tau-containing neurofibrillary tangles. Clinically, the hallmark of AD is cognitive decline, and the glutamatergic system, specifically the NMDA receptor (N-methyl-D-aspartate receptor, NMDAR), plays a major role in cognitive processes [1, 2]. NMDARs, which are essential for synaptic plasticity and memory formation, are highly concentrated in the human hippocampus [3, 4]. As such, loss of functional NMDAR and the resulting impairment in NMDA-evoked Long-Term Potentiation (LTP) is a central mechanism in AD that links the initial memory loss with the characteristic neurodegeneration of the hippocampus [5–7].

Studies in humans and in animal models have described multiple interactions between amyloid beta, phosphorylated tau and various glutamatergic transmission markers, underscoring the importance and complexity of the role played by NMDAR in AD [8–19]. Early work indicated loss of NMDAR measured by autoradiography in postmortem cortex and hippocampus of AD patients [20–22]. This was confirmed by our own autoradiographic study [23] demonstrating AD-related decreases in NMDAR density in hippocampus and entorhinal cortex, which tracks with disease severity as assessed by Braak staging. Conversely, several studies using different techniques found evidence for increased NMDAR protein and activation in early AD [16, 24, 25]. Reasons behind the inconsistent results are unknown, however, the above-mentioned studies did not examine the effects of sex or the relationship between regional NMDAR density and antemortem cognitive status across the AD spectrum. This is an important gap in the field since sex differences in hippocampal expression of NMDARs may contribute to the known sex differences in the cognitive trajectory of the disease, characterized by a female cognitive advantage in early disease stages followed by accelerated decline once pathology surpasses a critical threshold [26, 27].

Utilizing *in vitro* autoradiography with the specific NMDAR antagonist [^3^H]-MK801, we performed the first fully quantitative analysis of NMDAR density in the hippocampal formation, entorhinal cortex, and the parietal cortex of women and men who died with a clinical diagnosis of AD dementia, Mild Cognitive Impairment (MCI, the prodromal stage of AD), or cognitively normal (CN) controls. Furthermore, we examined the relationship between regional NMDAR density and antemortem cognitive function assessed by the Mini-Mental State Examination test (MMSE), and between NMDAR and tau density in women and men across the AD trajectory.

## 2. Results

### 2.1. Sample Characteristics

The characteristics of men and women in each diagnostic group are summarized in Table 1. There were no significant main effects of diagnosis, sex, or their interaction on postmortem delay or years of education (two-way ANOVAs, all p > 0.05). For age, a two-way ANOVA revealed a significant main effect of diagnosis (p = 0.0005), with no significant effect of sex (p = 0.91) or diagnosis × sex interaction (p = 0.995). Thus, the MCI group showed the highest mean age in both sexes (Table 1), although pairwise post-hoc comparisons did not reach significance. Within the AD group specifically, ApoE4 allele carriership was more frequent in men with AD compared to men in the CN and MCI groups (p < 0.01, Fisher’s exact test); while in women, ApoE4 carriership frequency did not differ significantly across the CN, MCI, and AD diagnostic groups.

**Table 1.** Characteristics of tissue donors by sex.

| Characteristics | Women (n = 76) |  |  | Men (n = 79) |  |  |
| --- | --- | --- | --- | --- | --- | --- |
|  | CN<br>(n = 25) | MCI<br>(n = 24) | AD<br>(n = 26) | CN<br>(n = 26) | MCI<br>(n = 26) | AD<br>(n = 27) |
| Age years (SD) | 83.1 (12.0) | 89.0 (6.6) | 83.3 (11.1) | 82.8 (7.2) | 89.0 (6.4) | 83.0 (7.1) |
| Education (SD) | 14.6 (2.6) | 14.9 (3.1) | 14.1 (2.7) | 15.3 (3.3) | 14.5 (2.6) | 14.7 (3.2) |
| Non-Hispanic white,<br>n (%) | 25 (100) | 23 (100)<sup>a</sup> | 24 (92) | 24(100)<sup>b</sup> | 26 (100) | 25 (100)<sup>*</sup> |
| APOE e4 carriers,<br>n (%) | 5 (20) | 5 (21) | 6 (24)<sup>a</sup> | 4 (15) | 8 (31) | 16 (59) |
| PMI hours (SD) | 3.6 (1.3) | 4.3 (2.4) | 3.3 (1.0) | 4.6 (2.3) | 4.0 (1.3) | 3.6 (1.5) |
| Global Cognition<br>(MMSE) | 28.2 (1.8) | 25.4 (3.1) | 14.6 (7.6)<sup>*</sup> | 27.9 (2.0) | 24.9 (2.7) | 14.6 (8.7)<sup>*</sup> |

Values are presented as mean, standard deviation (SD). LJInformation unavailable from one subject. LJInformation unavailable from two subjects. There were no significant effects of sex on any of the characteristics (ANOVA, p > 0.05). Significant effect of diagnosis on MMSE scores (p < 0.0001, two-way ANOVA); *Significantly different from CN and MCI groups; Tukey’s multiple comparisons test. Significant effect of diagnosis on age (p < 0.05, two-way ANOVA). CN = cognitively normal; MCI = mild cognitive impairment; AD = Alzheimer’s disease; PMI = postmortem interval; MMSE = Mini Mental State Examination.

### 2.2. The effect of diagnosis on NMDAR density is modulated by sex

The anatomical distribution of NMDARs in the hippocampus and cortex of CN individuals was similar to the known distribution of NMDARs in the human brain (Fig. 1, [23]), with highest densities in the CA1 (women: 10.8 ± 0.6; men: 9.8 ± 0.6) and dentate gyrus (DG; women: 10.8 ± 0.5; men: 9.6 ± 0.4) and lowest densities in cortical white matter (women: 1.0 ± 0.51; men: 0.15 ± 0.08).

**Figure. 1.**
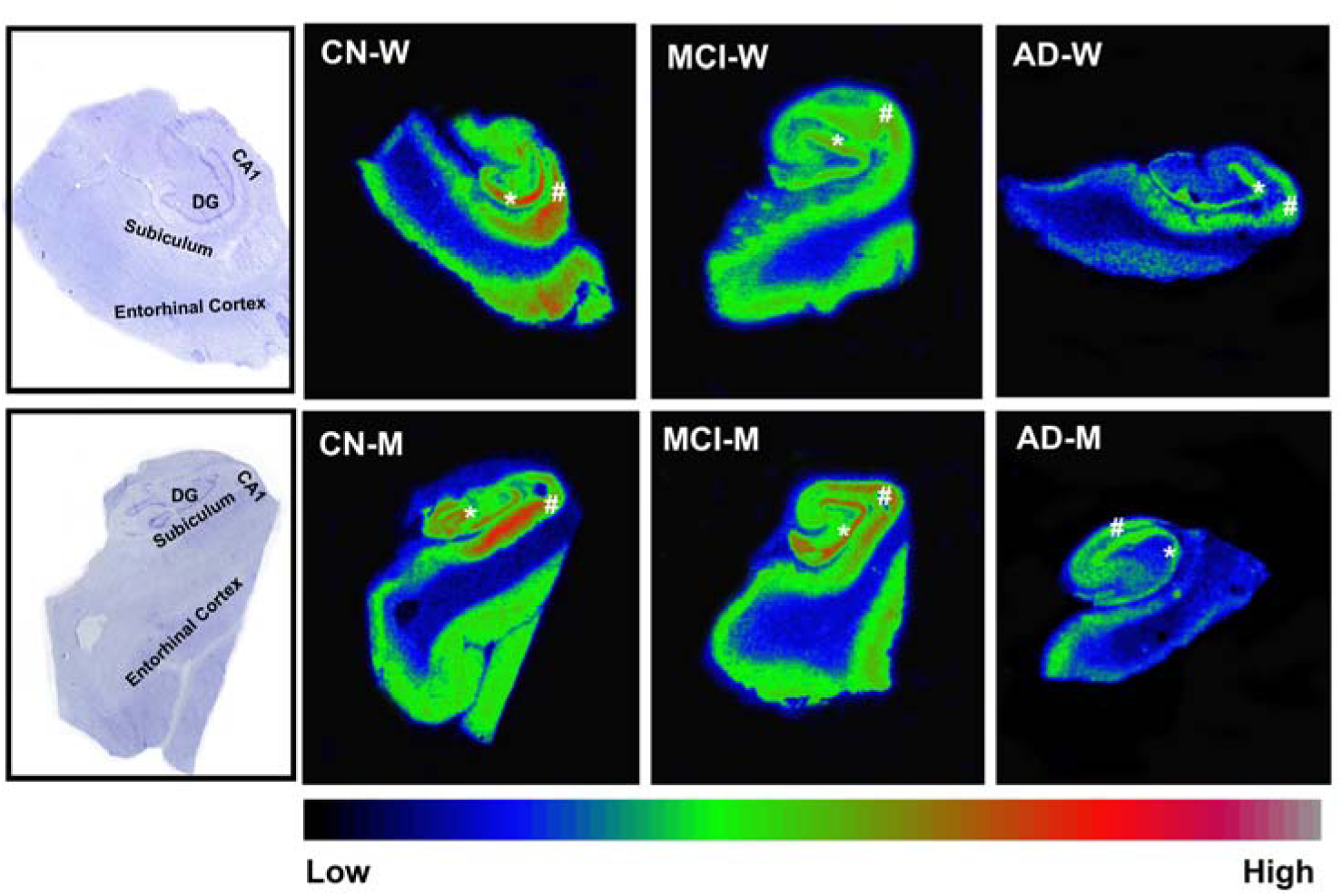
NMDA receptor density in the hippocampus and adjacent entorhinal cortex of women and men with antemortem clinical diagnosis of cognitively normal (CN), mild cognitive impairment (MCI), and Alzheimer’s disease (AD) dementia measured with [^3^H]-MK801 autoradiography. Verification of anatomy is illustrated in the leftmost panel, showing histological (cresyl violet) staining of consecutive sections from adjacent CN subjects, followed by representative pseudocolored autoradiograms of NMDA receptor binding labeled with [^3^H]-MK801 from the hippocampus and entorhinal cortex of a CN, MCI, and AD women (top) and men (bottom). Autoradiograms were pseudocolored using the Rainbow RGB lookup table (bottom bar); asterisks point to the DG, hash point to the CA1. Note highest NMDA receptor density observed in the CA1 and DG of CN subjects. Abbreviations: W, Woman; M, Man; CA1= cornu ammoni field 1 of the hippocampus; DG, dentate gyrus; CN, cognitively normal; MCI, mild cognitive impairment; AD, Alzheimer’s disease

Three-way ANCOVA (factors: region, diagnosis, and sex) of NMDAR density, with age and years of education as covariates, showed a significant effect of region (p < 0.001; DG > CA1 > subiculum (S) > entorhinal cortex (EC) > parietal cortex), a significant main effect of diagnosis (p = 0.045), and a highly significant sex × diagnosis interaction (p = 0.007). There was no significant main effect of sex (p = 0.447), no significant sex × region interaction (p = 0.838), no significant region × diagnosis interaction (p = 0.240), and no significant sex × region × diagnosis interaction (p = 0.968). Moreover, covariates age (p = 0.950) and years of education (p = 0.838) did not have a statistically significant effect on NMDAR density; the three-way ANCOVA was therefore re-run as a three-way ANOVA without covariates and yielded the same pattern of significant effects (data not shown).

To probe the significant sex × diagnosis interaction, we performed sex-stratified 2-way ANOVAs with region and diagnosis as fixed factors. A significant main effect of region and diagnosis on NMDAR density was observed in both sexes (region, p < 0.01; diagnosis, women, p = 0.01; men, p = 0.03); however, a sex-specific pattern in the diagnosis effect emerged, whereby women showed lower NMDAR density in both MCI and AD groups relative to CN (Fig. 2A), whereas men showed lower NMDAR density only in the AD group relative to the MCI and CN groups (Fig. 2B). This pattern reached statistical significance exclusively in the CA1 field of both men and women (Fig. 2A-B; women: AD<CN, p < 0.05; MCI < CN, p = 0.06, MCI = AD; men: MCI > AD, p < 0.05; CN > AD, p = 0.06, CN = MCI).

**Figure. 2.**
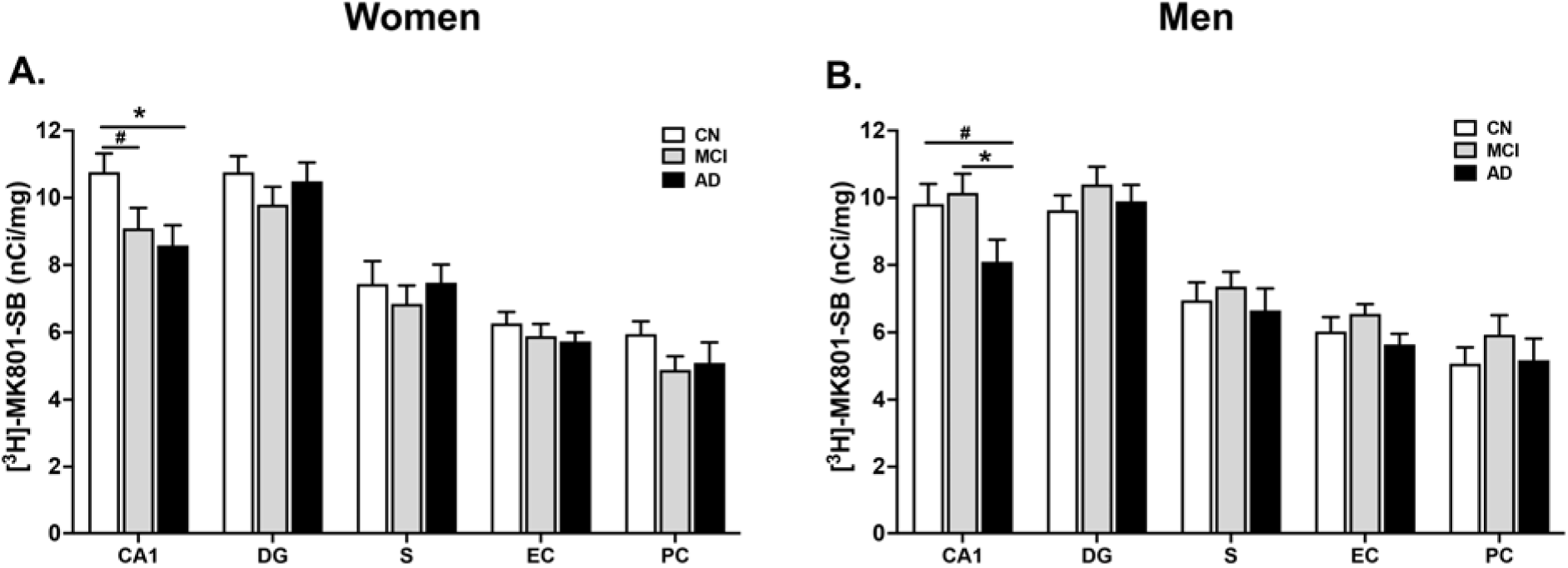
Diagnosis effects on NMDA receptor densities across hippocampal subregions and cortical regions are modulated by sex. Results of quantitative autoradiographic measurements of regional NMDA receptor density (expressed as specific binding of [^3^H]-MK801 in nCi/mg tissue) across diagnosis in women (**A**), and men (**B**). *p < 0.05, AD relative to CN in women or relative to MCI in men; #, approaching significance, p = 0.06. Results are from 2-way ANOVAs following a significant sex by diagnosis interaction (p<0.01) in 3-way ANCOVA with age and years of education as covariates. Bonferroni used for correction. Note a similar patten of a reduction in NMDA receptor density in both MCI and AD women relative to CN across regions, whereas in men NMDAR densities in MCI are maintained at levels that are not significantly different from CN. Data are presented as mean ± SEM. Abbreviations: CA1, cornu ammoni field 1 of the hippocampus; DG, dentate gyrus; S, subiculum; EC = entorhinal cortex; PC, parietal cortex; SB, specific binding; CN, cognitively normal; MCI, mild cognitive impairment; AD, Alzheimer’s disease.

### 2.3. The relationship between NMDAR density and cognitive scores is positive and significant in men but not women with AD

Next, we examined the relationship between NMDAR density and cognitive status as measured by the MMSE. Across sexes and diagnostic groups, we found a significant positive correlation between NMDAR density in the CA1 and MMSE scores (n = 124, r = 0.28, p < 0.01; Table 2). These correlations appeared to be driven by the low density/low performing men, confirmed when correlations were examined within sex and diagnostic groups: NMDAR levels in the CA1, DG, and EC were significantly and positively correlated with MMSE in AD men (CA1: n = 20, r = 0.67, p < 0.01; DG: n = 21, r = 0.64, p < 0.01; EC: n = 19, r = 0.50, p < 0.05; Table 2 and Fig. 3A-C), but there were no significant correlations between NMDAR density and MMSE in AD women or in men or women within the MCI and CN groups (Table 2 and Fig. 3D-F).

**Figure 3.**
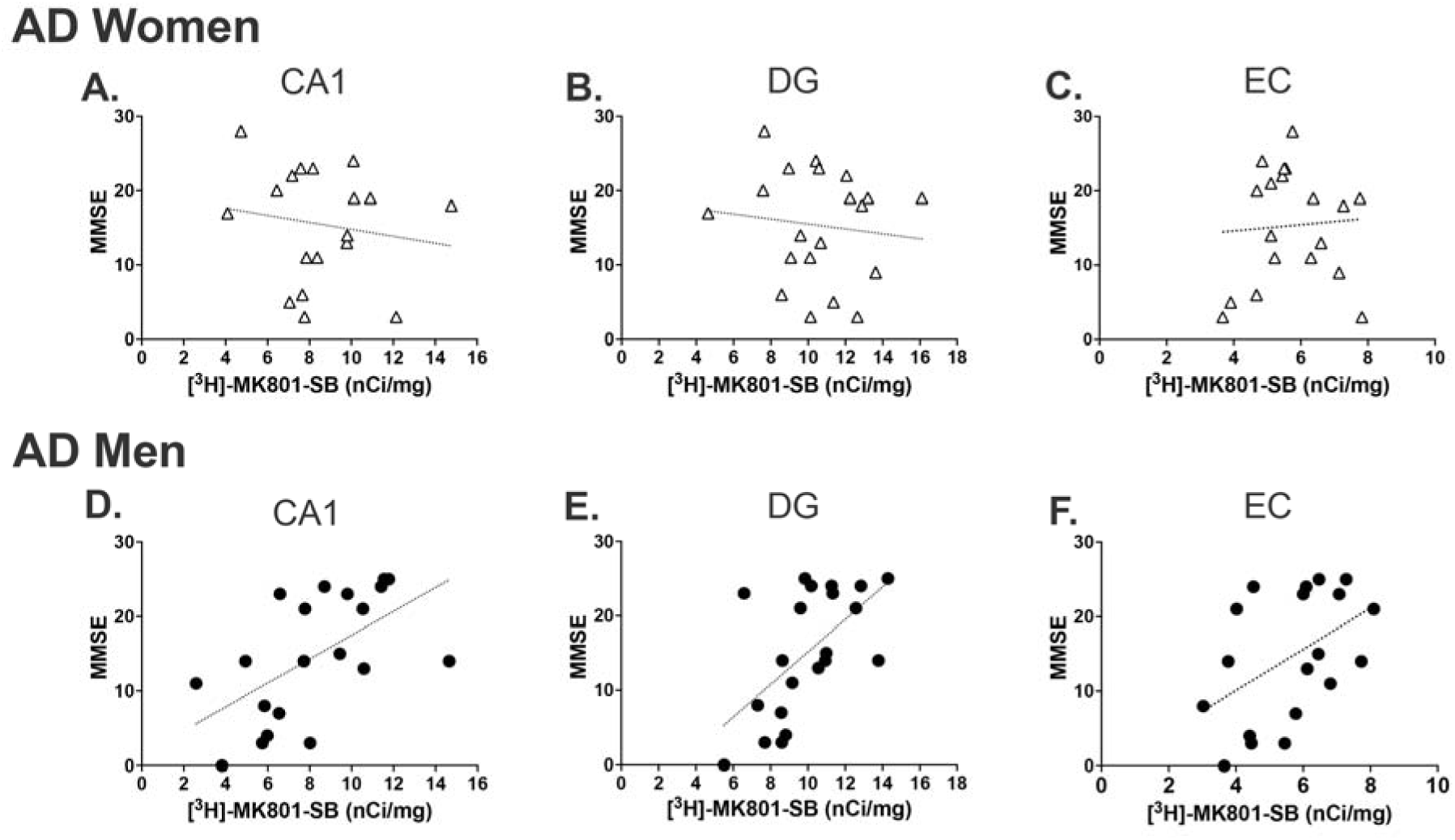
Significant positive correlations between MMSE scores and NMDAR density in men with Alzheimer’s disease. Note that statistically significant positive correlations between MMSE scores and NMDAR density in the CA1, DG and EC were observed in men but not in women with AD. (A) AD men, n = 20, r = 0.67, p < 0.01; (B) AD men, r = 0.64, n = 20, p < 0.01; (C) AD men, r = 0.50, n = 19, p < 0.05; (D) AD women, r = −0.32, n = 21, p > 0.05; (E) AD women, r = −0.15, n = 20, p > 0.05; (F) AD women, r = 0.04, n = 18, p > 0.05.

**Table 2.**
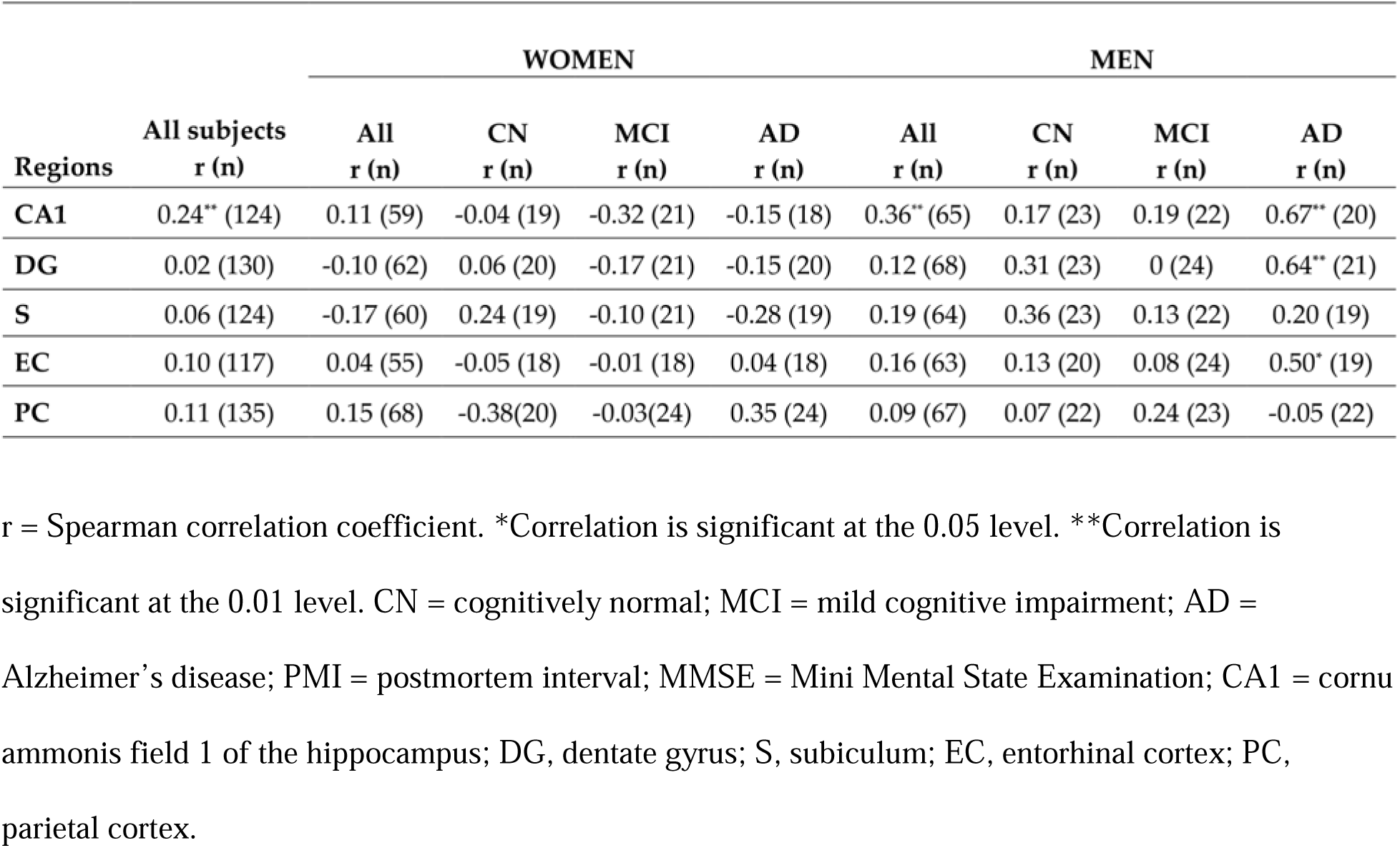
Results from Spearman’s correlations between NMDAR density and antemortem cognitive scores on the MMSE.

### 2.4 Tau density is elevated in AD and shows a sex-specific regional distribution

We sought to examine the relationship between NMDAR regional density and the core pathology of AD, namely density of tau deposits. To this end, we performed quantitative *in vitro* autoradiography with [^18^F]-T807 on consecutive sections from the same brain samples in which we measured NMDAR density. As expected, tau deposition was low to non-existent in the CN group, with low to moderate densities in the MCI group, and high density in the entorhinal cortex in the AD group (Figure. 4).

**Figure 4.**
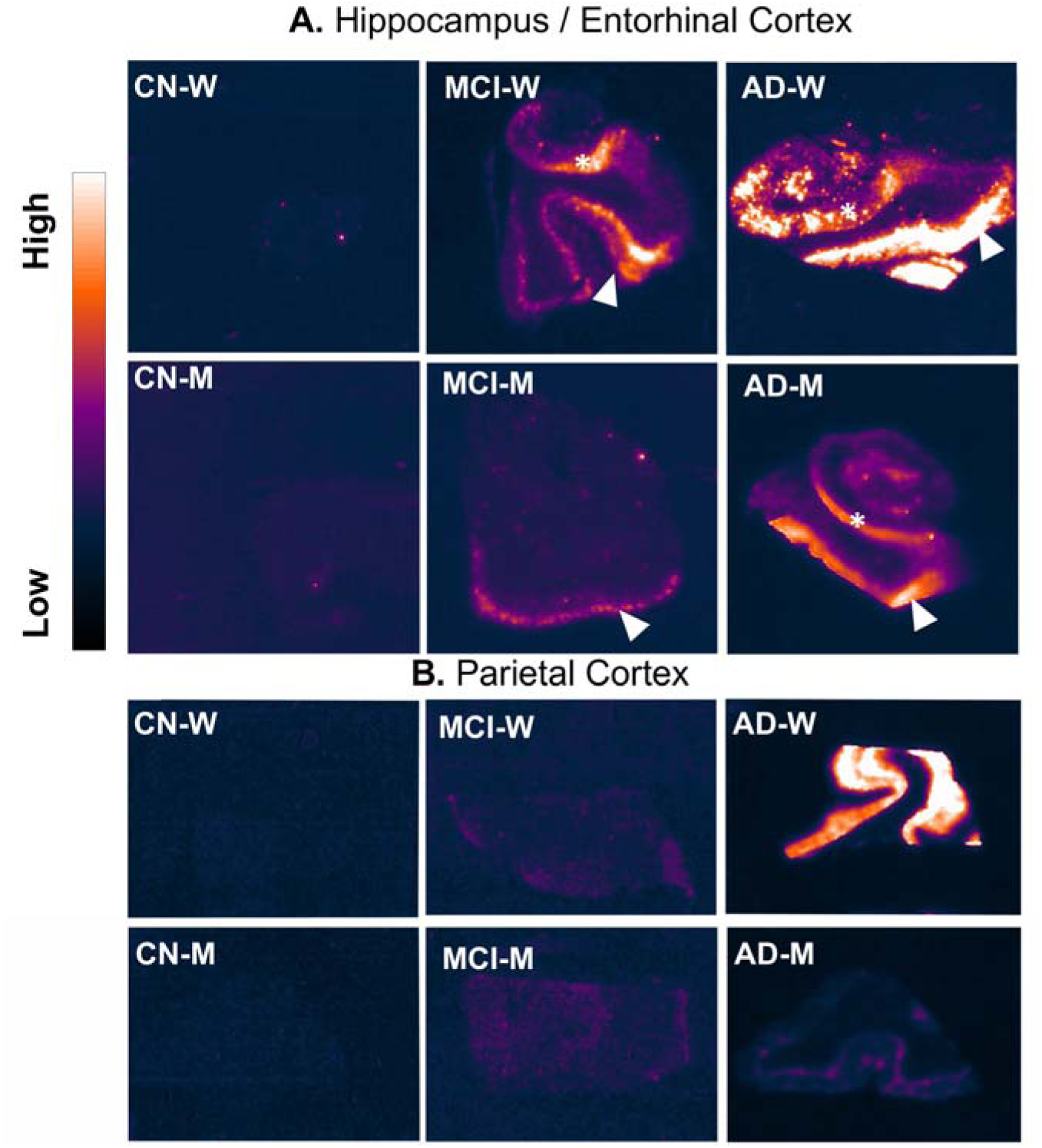
Densities of tau in the hippocampus, adjacent entorhinal cortex and the parietal cortex of CN, MCI, and AD women and men measured with [^18^F]-T807 autoradiography. Panels depict representative pseudo-colored autoradiograms of tau labeled with [^18^F]-T807 from the hippocampus and adjacent entorhinal cortex (panel A) and from the parietal cortex (panel B) of CN, MCI and AD subjects. Autoradiograms were pseudo-colored using the Gem lookup table (bar on the left). White arrows point to the entorhinal cortex and white asterisks point to the subiculum in MCI and AD subjects. Note the high tau density in the entorhinal cortex of AD subjects and in the parietal cortex of AD women.

A three-way ANCOVA was conducted to analyze the effects of diagnosis, sex, and region on cube-root transformed tau density, while controlling for age and years of education (covariates). There was a significant effect of age (p = 0.020) and of years of education (p = 0.005) on tau density; both covariates were therefore retained in the final model. After covariate adjustment, there was a significant main effect of sex (p < 0.001), diagnosis (p < 0.001), and regions (p < 0.001) on tau density. There were also significant two- and three-way interactions of sex x diagnosis (p < 0.001), diagnosis x region (p < 0.001), and sex x diagnosis x region (p = 0.034). The sex x region interaction did not reach significance (p = 0.110). Pairwise comparisons (Bonferroni-corrected) indicated that within the AD group, women showed higher tau density than men (p < 0.001; Figure. 4 and Figure. 5). Moreover, in women, tau density was significantly higher in the AD group relative to CN and MCI groups (p < 0.001; Fig. 5A), which did not differ from each other, while in men, tau density was significantly higher in the AD and in the MCI groups relative to CN (men: AD > CN, p < 0.001; MCI > CN, p = 0.03; AD > MCI, p < 0.001). Subsequent analyses stratified by sex revealed significant main effects of diagnosis and region in both sexes, but only women showed a significant diagnosis x region interaction (p < 0.001). The diagnosis x region interaction in women was driven by higher cortical tau density relative to hippocampal subregions within the AD group (tau density in AD women: EC > CA1, p = 0.008; EC > DG, p < 0.001; parietal cortex (PC) > CA1, p < 0.001; PC > DG, p < 0.001; PC > subiculum, p = 0.03; EC = PC). Figure 5 shows results stratified by sex with Bonferroni-corrected pairwise comparisons for each region.

**Fig. 5.**
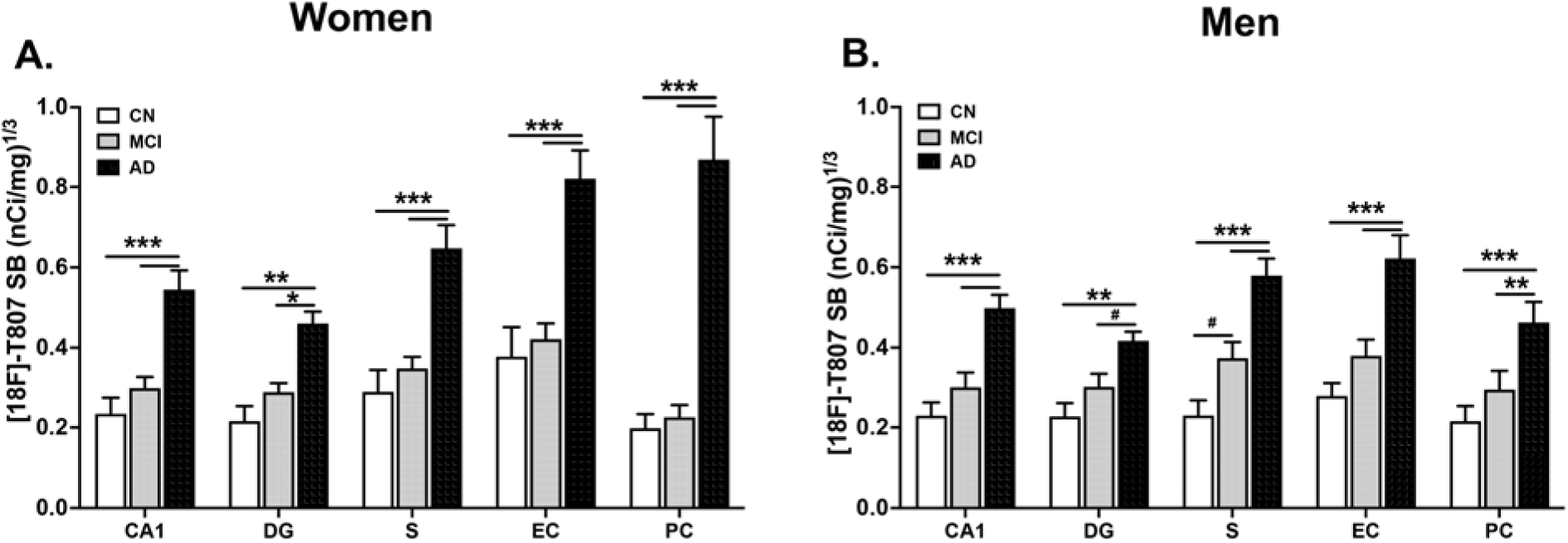
Tau densities are significantly elevated in hippocampal subregions and cortical regions of men and women with AD. Graphs depict quantitative autoradiographic measurements of tau density (expressed as specific binding of [18F]-T807 in nCi/mg tissue) across diagnosis in women (A) and men (B). ***p < 0.001, **p < 0.01, *p < 0.05, AD relative to controls or MCI, #, approaching significance, p = 0.06 - 0.07. Two-way ANCOVA with age and years of education as covariates followed by Bonferroni correction. Data are presented as mean ± SEM. Abbreviations: CA1, cornu ammoni field 1 of the hippocampus; DG, dentate gyrus; S, subiculum; EC = entorhinal cortex; PC, parietal cortex; SB, specific binding; CN, cognitively normal; MCI, mild cognitive impairment; AD, Alzheimer’s disease.

### 2.5 NMDAR density is negatively correlated with tau density in AD men

Analysis segregated by sex and by diagnostic group revealed negative correlations between NMDAR and tau density in men with AD (Table 3). The highest and most significant negative correlation was noted in the EC of men with AD (n = 19, r = -0.60, p < 0.01), followed by the CA1 (n = 20, r = -0.44, p = 0.05) and the DG (n = 20, r = -0.34, p = 0.10), where negative correlation between NMDAR and Tau density approached but did not reached significance (Table 3). On the other hand, in AD women negative correlations between NMDAR and tau density approached significance only in the PC (n = 23, r = -0.38, p = 0.08).

**Table 3.**
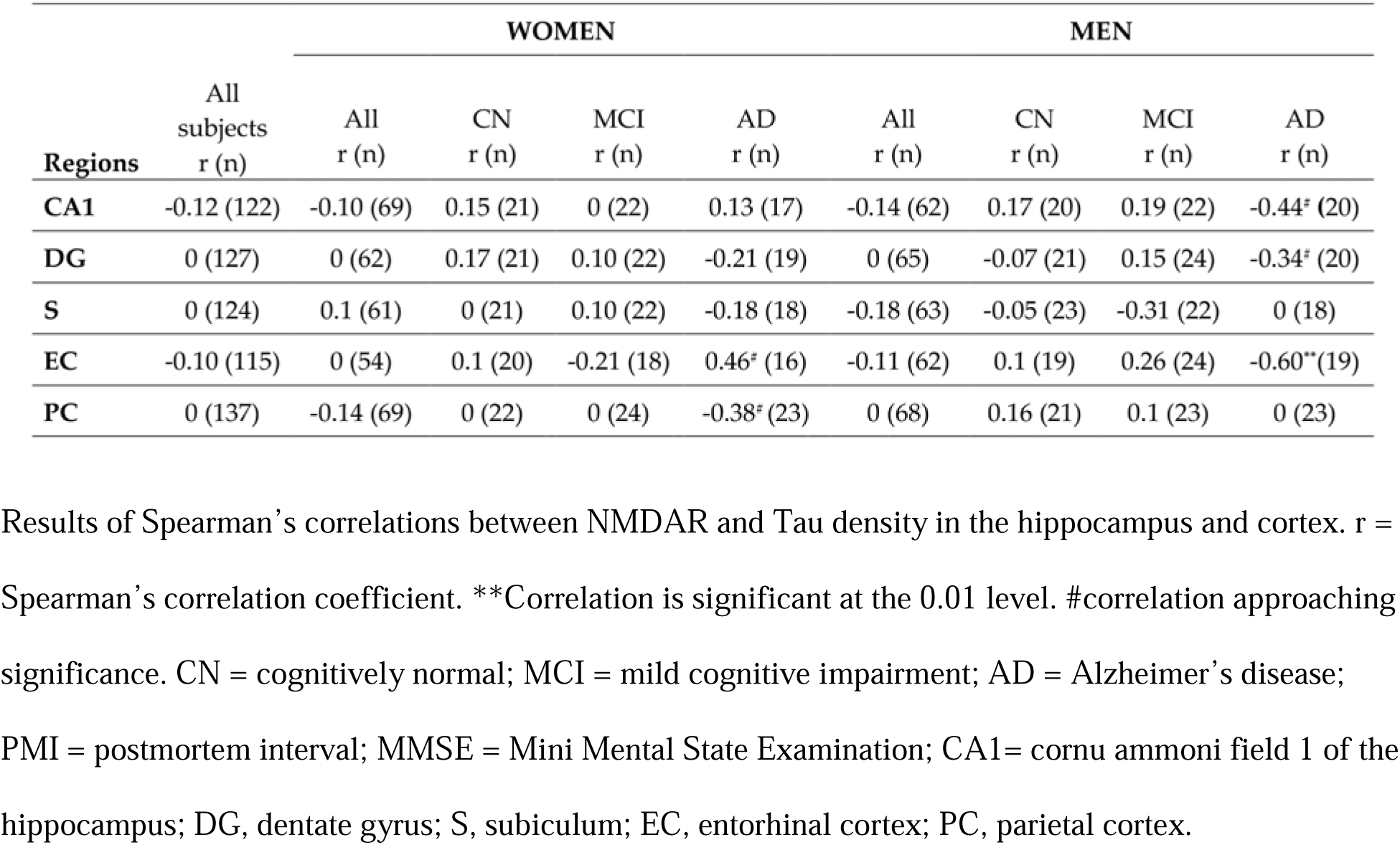
Spearman correlations between regional NMDAR density and tau density, stratified by sex and diagnosis.

Notably, side by side comparison of tau and NMDAR autoradiograms from the same individuals offer an explanation to the correlation findings since, unlike NMDAR density, tau deposition appears as puncta and clusters. As can be seen in Figure 6, clusters of very high tau density coincide with locally lower NMDAR density within hippocampal and cortical regions, whereas areas with low/punctate tau deposition show relatively higher NMDAR density.

**Figure. 6.**
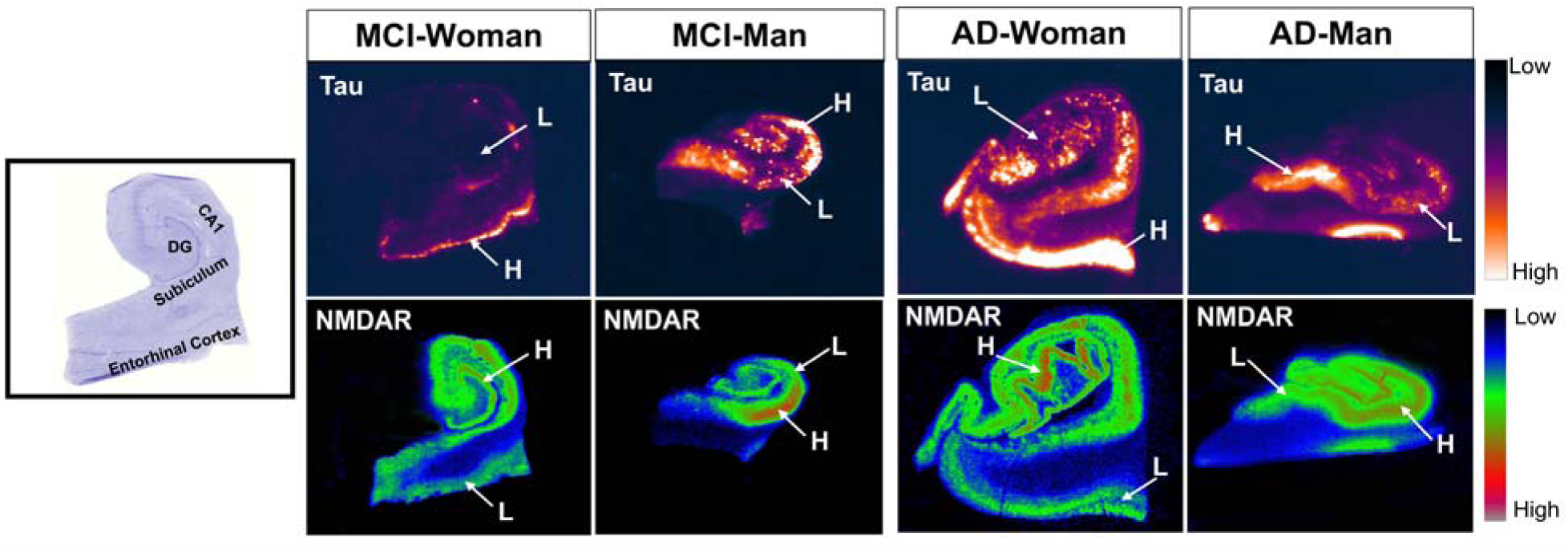
Clusters of very high tau density coincides with locally lower NMDA receptor density within hippocampal and cortical regions. Top panels depict representative pseudo-colored autoradiograms of tau labeled with [^18^F]-T807 from the hippocampus and adjacent entorhinal cortex of female and male MCI subjects and female and male AD subjects. Bottom panels show representative pseudo-colored autoradiograms of NMDA receptor labeled with [^3^H]-MK801 from consecutive sections of the hippocampus and entorhinal cortex from the same subjects. Arrows point to areas of high (H) tau density coinciding with areas of low (L) NMDA receptor density, whereas low (L) tau density coincides with areas of high (H) NMDA receptor density.

## 3. Discussion

In this quantitative autoradiographic study of NMDAR in hippocampal and cortical regions across the AD trajectory, we show that a clinical diagnosis of AD is associated with reductions in NMDA receptor density in the hippocampus of both women and men, with the most pronounced loss in the CA1 field. Among subjects with a diagnosis of MCI, women, but not men, exhibit lower NMDAR density relative to age-matched controls. Lower NMDAR densities are associated with poor antemortem cognitive performance as measured by the MMSE. Moreover, higher tau density is associated with lower NMDAR density, with significant associations localized to the entorhinal cortex of AD men.

The fate of NMDAR in the brain of subjects with AD has been the subject of a number of previous studies reporting changes in NMDARs. Using L-[^3^H]-glutamate binding to sections from the hippocampal formation of six patients who died from dementia of the Alzheimer’s type (DAT) and six patients without DAT, Greenamyre et al. [20] found marked reductions in total [^3^H]-glutamate binding in all regions of hippocampus and adjacent parahippocampal cortex in DAT brains as compared to controls. When subtypes of excitatory amino acid receptors were assayed, it was found that binding to the N-methyl-D-aspartate (NMDA)-sensitive receptor was most markedly reduced, with the greatest loss found in stratum moleculare and stratum pyramidale of CA1. We replicated and extended these findings in a subsequent autoradiographic study showing reduced NMDAR binding in the CA1 of subjects who died with AD which tracked disease severity by Braak stage and was negatively correlated with neuroinflammation [23]. In a recent study, lower levels of synaptic NMDAR subtypes GluN2B and GluN2A were found in AD synaptic membrane fractions, whereas extrasynaptic GluN2B and GluN1 levels were significantly higher in AD [25]. This apparent contradiction may suggest that both decreases and increases in NMDAR density may occur, either sequentially or in parallel, in different subcellular locations and stages of disease progression and may also be modulated by sex [28].

To elaborate, there is a near-consensus that synaptic loss is an early event in AD [29–32]. Synaptic degeneration and the ensuing loss of contacts results in sustained decreases in NMDAR stimulation, impairing synaptic plasticity in the form of LTP and LTD, both necessary for learning and memory [2, 6, 14]. Conversely, sustained reductions in synaptic transmission – through persistent decreases in NMDAR stimulation – can also drive compensatory upregulation of NMDARs. This has been demonstrated by *in vitro* and *in vivo* radioligand-binding studies in which a reduction in NMDAR stimulation in the form of chronic antagonist treatment leads to upregulation of NMDAR densities [33–36]. In fact, postmortem studies have shown up-regulation of NMDAR subunits (ie. GluN1, GluN2B) in the early stages (Braak II) of AD [16]. Other studies suggest that increased NMDAR density early in AD may reflect up-regulation of extrasynaptic receptors, whose low but continuous stimulation contributes to synaptic impairment [33]. Hence, the changes in NMDAR densities reported in our study likely may reflect both a compensatory mechanism observed in the early stages of AD, as well as altered function of synaptic / extrasynaptic NMDARs that initiate hyperactivation and eventual excitotoxicity, with the end result being loss of NMDAR positive neurons tipping the balance towards lower receptor density in advanced AD.

With regard to sex differences in NMDAR density, we uncovered a divergent trajectory in the loss of NMDAR among women and men with MCI: in the CA1, MCI women showed NMDAR density comparable to AD levels and lower than age-matched CN, whereas MCI men showed NMDAR levels slightly higher than age-matched CN and significantly higher than AD men. Mild cognitive impairment is considered a transitional stage from healthy aging to dementia, in which individuals – while still capable of performing activities of daily living – experience subtle but measurable cognitive deficits [34]. The fact that MCI women and not MCI men are already showing loss of NMDARs is consistent with studies demonstrating that compared to men, women experienced greater pathological burden at earlier stages of the disease [35–39]. In fact, women decline faster from MCI to AD [40, 41] and demonstrate heightened sensitivity to tau deposition [42, 43] and neuroinflammation [44]. In a recent longitudinal study by our group, we show that in the MCI stage, women exhibit a significantly steeper memory decline associated with increasing CSF pTau181/Ab42 ratios compared to men [27]. Therefore, the decrease in NMDAR density observed in MCI women might be another biological mechanism contributing to their steeper cognitive decline relative to men.

Surprisingly, while initial analyses that included all subjects revealed significant positive correlations between antemortem MMSE scores and CA1 NMDAR density, subsequent analyses stratified by sex and diagnosis showed that AD men — particularly the lower cognitive performers — drove this positive relationship. Although this pattern initially appears counter-intuitive given that women bear a disproportionately greater AD burden, it is consistent with a growing literature documenting sex-specific coupling between pathology and cognition. Women appear to tolerate more tau pathology before verbal-memory decline emerges [45], and in a multicenter memory clinic cohort spanning the full clinical spectrum, the association between tau load and cognitive performance was significantly stronger in men than in women, despite women showing greater tau burden overall [46]. By analogy, our NMDAR–MMSE finding suggests that in men — who may lack the compensatory or reserve mechanisms invoked by women — molecular measures of synaptic integrity map more directly onto cognitive performance.

Whether this reflects diminished reserve or differential downstream sensitivity to receptor loss remains an important question for future study. Notably, the persistence of this male-specific coupling into the AD-dementia stage — rather than being confined to earlier, preclinical stages — suggests that sex differences in pathology-cognition coupling may not be strictly stage-limited, and longitudinal designs tracking both sexes across the full AD continuum will be needed to disentangle stage-specific from sex-specific contributions to this pattern.

In both sexes, a statistically significant AD-related decrease in NMDAR density was observed only in the CA1 region. This CA1 vulnerability is in line with a substantial body of evidence from *in vivo* imaging and postmortem histological studies of AD, which consistently show selective vulnerability of the CA1 to volume loss and neuronal death relative to other hippocampal subfields [47–51]. In addition, ample literature demonstrates selective vulnerability of the CA1 in response to insults including excitotoxicity and ischemia [49, 52], and offers multiple explanations for this phenomenon [53, 54]. Moreover, in women and men with AD, we observed high tau deposition in the entorhinal cortex, which is the first cortical region to show significant tau deposition (Braak stages I-II; [55]). The CA1 region has direct, monosynaptic excitatory input from the EC [56–59]. This direct input is thought to create a neuronal pathway for the propagation of tau pathology from EC to CA1, with tau toxicity driving early neuronal death and neurodegeneration [60]. Because the EC serves as the major gateway for cortical input into the hippocampus, tau-induced damage to EC projection neurons deprives CA1 of its principal excitatory drive, contributing to CA1 functional isolation, neurodegeneration, and cell death.

In agreement with previous findings, we showed AD-related increases in tau density in all regions examined, with AD women demonstrating higher tau density relative to AD men [42–44]. Moreover, in women, there was a significant region-by-diagnosis interaction caused by prominent increases in the EC, the site of initial cortical tau deposition in AD, and in the PC, a region that shows tau deposition only at more advanced stages of the disease. In vitro and in vivo studies have shown complex interactions between tau and NMDARs, with studies suggesting that under pathological states, tau accumulation impairs synaptic plasticity through JAK2/STAT1-induced suppression of NMDAR expression [14]. However, despite the highly significant tau deposition in our AD subjects, tau density was significantly associated with decreased NMDAR density only in the EC of AD men, whereas in AD women, a negative association between tau and NMDAR density approached significance only in the parietal cortex. Taken together with the male specific correlation between CA1 NMDAR density and MMSE scores in AD, this pattern suggests that in men, NMDAR loss may be more directly linked to tau related neurodegeneration and consequent cognitive impairment, whereas in women these relationships may be more multifactorial or partly uncoupled from NMDAR density, potentially reflecting a broader set of pathological processes contributing to cognitive decline. We speculate that the lack of significant negative associations between tau and NMDAR density in additional regions is due to qualitative differences in the density patterns of these two markers within regions, whereby punctate tau deposition is often associated with preserved NMDAR density, while dense and confluent tau accumulation coincides with locally reduced NMDAR density. Nevertheless, the fact that negative correlations were found in regions with the highest tau density in women (PC) and in men (EC) is consistent with previous reports in which hippocampal protein levels and mRNA expression for specific NMDAR subunits, such as GluN1/GluN2B, are reduced with increasing AD-related neuropathology (Braak stage; [3]).

## Conclusion

NMDA receptor density is strongly modulated by sex, region and AD disease stage. Antemortem cognitive scores in men were more sensitive to loss of NMDAR. These findings may contribute to the understanding of the known sex differences in AD prevalence, presentation and progression.

## 4. Materials and Methods

### 4.1. Sample characteristics and handling

The current study used 10-micron thick sections of fresh-frozen samples of the hippocampus/entorhinal cortex and the parietal cortex from 154 donors equally divided among men and women and the three diagnostic groups (Control, MCI and AD). One hundred and seventeen samples (18-20 per sex / diagnosis) were obtained from the the Arizona Study of Aging and Neurodegenerative Disorders and Brain and Body Donation Program [61] and 37 samples (6-7 per sex / diagnosis) from the University of Washington BioRepository and Integrated Neuropathology (BRaIN) repository. Inclusion criteria were complete neuropathological examination (e.g., Braak staging, Thal amyloid staging) and a comprehensive antemortem neurocognitive evaluation that indicated a clinical diagnosis of either cognitively normal, MCI or AD dementia as recently described [44]. For BRaIN subjects, cognitive evaluations were required to be within 18 months of death; for BBDP subjects, diagnoses were based on the closest available evaluation from the NACC Uniform Data Set battery. Pathological data confirmed the presence of AD dementia or its absence for control and MCI groups, although control or MCI subjects could have some degree of incidental pathology. Cases that were given a neuropathological diagnosis of a neurodegenerative disease other than AD were excluded. For most subjects, ApoE genotype was also provided (Table 1). Post mortem interval (PMI) was less than 12 hours for all samples (Table 1). Upon reception, samples were stored in a -80°C freezer until use. All frozen samples were cryo-sectioned (Leica cm 1950) at -20°C (10 μm) in 10 consecutive series and thaw- mounted onto positively charged glass slides (Fisher Scientific).

### 4.2. In vitro quantitative autoradiography

The labelling of NMDAR on tissue sections was done as described previously [23]. Briefly, on the day of the assay, consecutive slides from each subject were removed from the −80 °C freezer and allowed to reach room temperature. After a 30-minute prewash in 50 mM Tris-acetate buffer (pH 7.4), the sections were incubated for 3 hours at room temperature in 50 mM Tris-acetate buffer at pH 7.4 containing 5 nM [^3^H]-MK801 (a selective non-competitive NMDAR antagonist; specific activity 27.5 Ci/mmol, PerkinElmer Life Sciences, Waltham, MA, USA), 30 µM glutamate, and 10 µM glycine. Nonspecific binding was determined in the presence of 10 µM unlabeled MK801. At the end of the incubation, the sections were dipped for 5 seconds in ice-cold buffer and then washed for 90 minutes in cold, fresh buffer, followed by a dip in ice-cold double-distilled water. Sections were then dried on a slide warmer at 60 °C and apposed to low-energy radiation sensitive film (BluBlot HS, Chemglass) for 8 weeks, alongside calibrated tritium microscales (Amersham Pharmacia Biotech, Piscataway, NJ, USA). Films were developed in Kodak D-19, fixed, and dried. To assess tau deposits, autoradiography was performed with [^18^F]-T807 obtained from the Stony Brook University PET radiochemistry laboratory. Sections were transferred to a bath containing ∼2 nM [^18^F]-T807 in 10 mM PBS. After incubation for 60 minutes, slides were washed in 10 mM PBS for 1 minute, 70% ethanol / 30% PBS for 2 minutes, 30% ethanol / 70% PBS for 1 minute, and lastly in PBS for 1 minute [48]. Slides were then air dried, apposed to film for 1 hour and developed as described above. All sections were cresyl violet stained for histology after film development.

### 4.3. Quantitative image analysis

Developed films were scanned using a flatbed scanner (2,400 dpi, 256 gray levels, in neutral conditions—no highlights, shadows, midtones, gamma or sharpen manipulations), digitized and saved in 8-bit TIFF format. Quantitative image analyses were performed using the FIJI (NIH Image) software. Regions of interest (ROIs) in hippocampus samples were identified in reference to the histological staining and an atlas of the human hippocampus [65] and included the cornu ammonis field 1 (CA1), dentate gyrus (DG), subiculum (S), entorhinal cortex (EC) and parietal cortex (PC). ROIs were manually drawn on the autoradiograms around the whole extent of each region on all sections where the region could be clearly distinguished on the histologically stained section. Nonspecific binding was subtracted from total binding to generate specific binding values for statistical analysis.

Image analysis of autoradiograms and statistical analysis of the results from each experiment were performed by an investigator blinded to sex and diagnosis and then decoded by the senior investigators (M.A.M. and A.B.).

### 4.4. Statistical Analysis

Prior to modeling, distributional assumptions were examined using skewness and kurtosis values as well as the Shapiro–Wilk test. Tau density values were cube-root transformed to meet assumptions of normality. Homogeneity of variance was tested for all continuous variables using Levene’s test. Demographic data were analyzed using a two-way ANOVA with sex and diagnosis as factors. ApoE genotype was dichotomized into Apoε4 carriers and non-carriers, and Fisher’s exact tests were used to examine diagnosis effects.

For each autoradiographic marker (NMDAR density, tau density), the effects of diagnosis, sex and region were evaluated by a three-way ANCOVA with diagnosis (CN, MVI, AD), biological sex and region (CA1, DG, subiculum, EC, PC) as fixed factors, and age at death and years of education as candidate covariates. Covariates were initially evaluated in a full model and retained in the final model only when they exerted a significant effect (p < 0.05) on the dependent variable; when neither covariate reached significance, a three-way ANOVA without covariates was used. Under this rule, age and years of education were retained for tau density but not for NMDAR density. When a significant sex x diagnosis or sex x diagnosis x region interaction was detected, sex-stratified two-way ANOVA (or ANCOVA as appropriate) was performed with diagnosis and region as fixed factors. Pairwise post-hoc comparisons were corrected with the Bonferroni method.

Given the modest within-group sample sizes (n = 18-23 per sex x diagnosis cell) and the restricted, ceiling-affected distribution of MMSE scores, associations between regional NMDAR density and MMSE, and between regional NMDAR and tau density, were assessed with Spearman’s rank correlation.

Image analyses of autoradiograms and preliminary statistical analyses were performed by an investigator blinded to sex and diagnosis; group codes were revealed only for final interpretation by the senior investigators (MAM and AB).

All statistical analyses were performed using SPSS Statistics version 28 (IBM Corp., Armonk, NY); graphs were prepared with GraphPad Prism 9 (GraphPad Software, San Diego, CA). Statistical significance was set at α = 0.05. Data are presented as mean ± SEM unless otherwise noted.

## Author Contributions

Conceptualization, A.B., M.A.M., and E.E.S.; methodology, S.M., J.D.; validation, M.A.M., A.B., and S.M.; investigation, S.M., V.C., A.N., J.D.; resources, G.E.S., T.G.B., and M.A.M.; data curation, M.A.M.; writing—original draft preparation, M.A.M. and A.B.; writing—review and editing, E.E.S.; visualization, M.A.M.; supervision, M.A.M. and A.B.; project administration, M.A.M.; funding acquisition, E.E.S. All authors have read and agreed to the published version of the manuscript.

## Institutional Review Board Statement

This work was done at Stony Brook University using de-identified human tissue samples collected under approved protocols by established tissue repositories, namely the University of Washington (UW) BioRepository and Integrated Neuropathology laboratory (BRaIN; https://dlmp.uw.edu/research-labs/keene/BRaIN-lab) and the Arizona Study of Aging and Neurodegenerative Disorders and Brain and Body Donation Program (BBDP; www.brainandbodydonationprogram.org) after approval by their respective institutional review boards.

## Informed Consent Statement

All participants provided written informed consent, which was approved by the institutional review boards from the University of Washington or Banner Sun Health Research Institute.

## Data Availability Statement

The raw data supporting the conclusions of this article will be made available by the authors on request.

## Abbreviations

AD: Alzheimer’s disease
CA1: cornu ammonis field 1
CN: cognitively normal
DG: dentate gyrus
EC: entorhinal cortex
LTP: long-term potentiation
MCI: mild cognitive impairment
MMSE: Mini Mental State Examination
NMDAR: N-methyl-D-aspartate receptor
PC: parietal cortex
PMI: postmortem interval
S: subiculum.

